# Machine intelligence identifies soluble TNFa as a therapeutic target for spinal cord injury

**DOI:** 10.1101/2020.07.22.216572

**Authors:** JR Huie, AR Ferguson, N Kyritsis, J Z Pan, K-A Irvine, JL Nielson, PG Schupp, MC Oldham, JC Gensel, A Lin, MR Segal, RR Ratan, JC Bresnahan, MS Beattie

**Author notes:** Correspondence should be addressed to Adam R. Ferguson and Michael S. Beattie, Brain and Spinal Injury Center (BASIC), Department of Neurological Surgery, University of California San Francisco, San Francisco, CA. Equal contribution (JR Huie, AR Ferguson).

## Abstract

Traumatic spinal cord injury (SCI) produces a complex syndrome that is expressed across multiple endpoints ranging from molecular and cellular changes to functional behavioral deficits. Effective therapeutic strategies for CNS injury are therefore likely to manifest multi-factorial effects across a broad range of biological and functional outcome measures. Thus, multivariate analytic approaches are needed to capture the linkage between biological and neurobehavioral outcomes. Injury-induced neuroinflammation (NI) presents a particularly challenging therapeutic target, since NI is involved in both degeneration and repair^1,2^. Here, we used big-data integration and large-scale analytics to examine a large dataset of preclinical efficacy tests combining 5 different blinded, fully counter-balanced treatment trials for different acute anti-inflammatory treatments for cervical spinal cord injury in rats. Multi-dimensional discovery, using topological data analysis^3^ (TDA) and principal components analysis (PCA) revealed that only one showed consistent multidimensional syndromic benefit: intrathecal application of recombinant soluble TNFα receptor 1 (sTNFR1), which showed an inverse-U dose response efficacy. Using the optimal acute dose, we showed that clinically-relevant 90 min delayed treatment profoundly affected multiple biological indices of NI in the first 48 hrs after injury, including reduction in pro-inflammatory cytokines and gene expression of a coherent complex of acute inflammatory mediators and receptors. Further, a 90 min delayed bolus dose of sTNFR1 reduced the expression of NI markers in the chronic perilesional spinal cord, and consistently improved neurological function over 6 weeks post SCI. These results provide validation of a novel strategy for precision preclinical drug discovery that is likely to improve translation in the difficult landscape of CNS trauma, and confirm the importance of TNFα signaling as a therapeutic target.

---

Numerous therapeutic targets have been identified for SCI, including treatments aimed at blunting or modulating the post-injury cascade of neuroinflammation^1^. But aspects of neuroinflammation are known to also initiate repair^4–7^. This multi-faceted aspect of neuroinflammation may be responsible for the difficulty in finding clean targets for therapeutics in this space, and for limited reproducibility. It is likely that this complex target responds differently to drugs affecting different aspects of the cascade, as well as to different drug doses. For example, tumor necrosis factor alpha (TNFα), one of the major pro-inflammatory mediators in this cascade, can have divergent effects at different doses due to differences in activation of its two canonical receptors, TNFR1 and TNFR2 (*TNFRSF1A and 1B*) ^8^. Minocycline, methylprednisolone, and other ‘antiinflammatory’ therapies also have pleiotropic effects on multiple signaling pathways, and their efficacy in preclinical treatments for SCI has been variable^9–13^. The complex nature of these drugs’ actions limits reproducibility of these findings, threatening the predictive potential of preclinical SCI research^14–16^. In the complex tissue microenvironment of the injured CNS, drugs with specific mechanisms of action often lack the breadth of efficacy required to improve behavioral function. Further, the arbitrary selection of a single *a priori* primary outcome metric may lead to a narrow and often-inadequate view of complex disease syndromes where multiple factors contribute to pathogenesis. Focusing on a single univariate outcome metric can thus lead to inaccurate assessments, wasted resources, and failed clinical trials^17^. Improved data integration and multivariate analytics offer opportunities to improve efficacy-testing^18–21^ and improve precision medicine for CNS trauma^22^.

Here, we used large-scale data-driven discovery techniques to extract syndromic information from complex SCI outcomes data across a series of experiments on neuroinflammatory targets. These data were sequentially obtained in a series of five independent studies that tested promising drugs with reputed anti-inflammatory properties (see **Supplemental Table 1**). We reasoned that combining these studies into an omnibus analytic would provide useful multivariate outcome measures that would leverage variability of the combined large-n dataset to optimize detection of potential treatments and targets. We first performed multidimensional data-curation/integration to incorporate ensemble information from preclinical health records, histology, and behavioral function from 5 different unpublished, blinded preclinical neuroinflammation trials performed over 10 years in our laboratories^23^. We then subjected this high-content data to topological data analysis (TDA) for data-driven discovery, pattern-detection, and dimensionality-reduction^21,24^ (**Fig. 1**). This approach was used in a prior *post hoc* analysis of historic VISION-SCI data derived from the multicenter animal spinal cord injury study (MASCIS, see^25^) and allowed us to discover latent predictors of outcome that characterize the entire SCI syndrome as an integrated data system. In that study, we discovered potent predictors of outcome related to physiological measures during SCI induction, and less predictive, but significant, drug effects that had not been found in univariate analyses. When applying TDA to our integrated neuroinflammation dataset, results revealed that only one of the anti-inflammatory interventions tested, intrathecal delivery of a soluble TNF receptor 1 (sTNFR1), yielded a consistent multidimensional benefit across the SCI syndromic space characterized by the TDA network topology.

**Figure 1.**
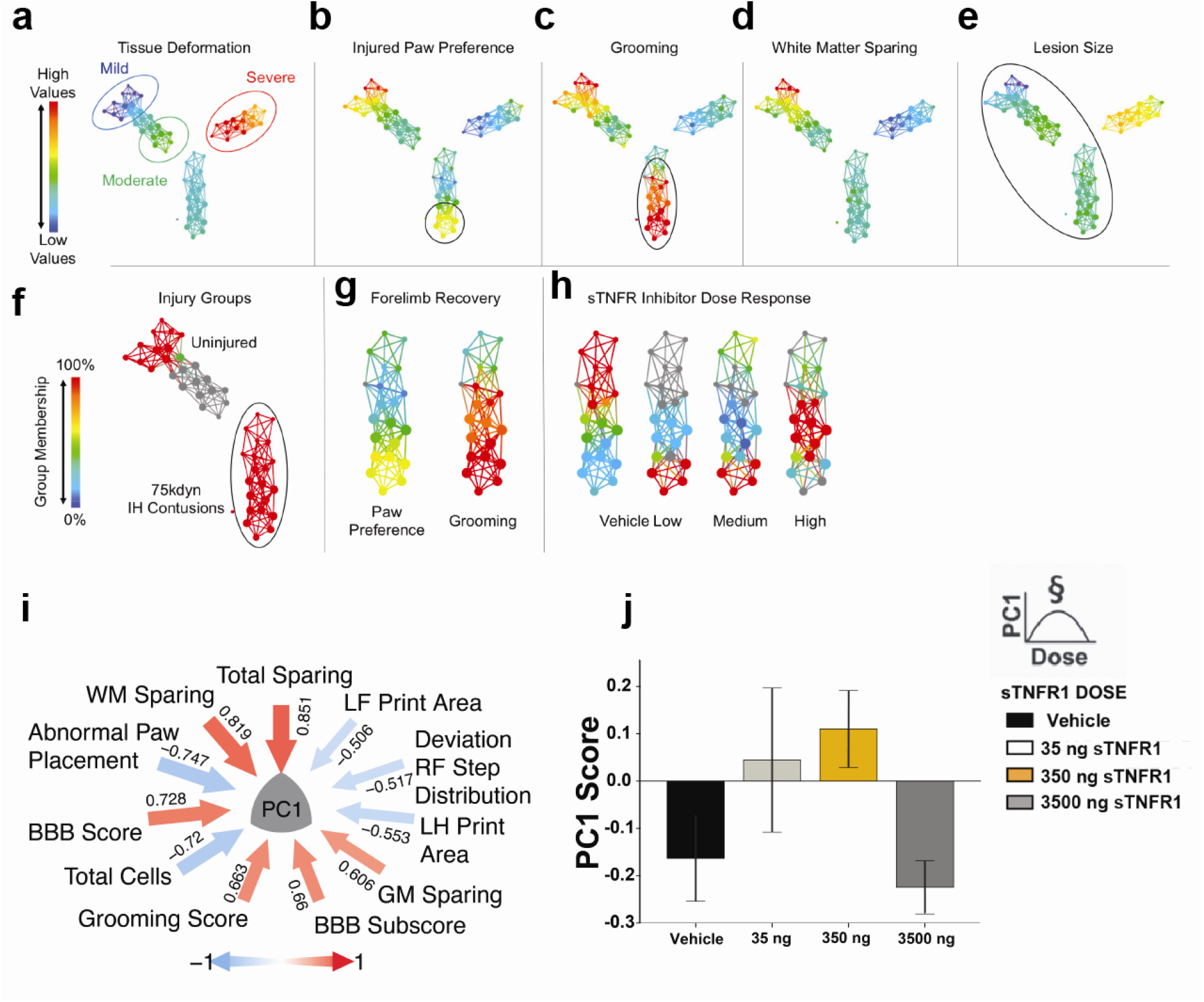
Topological Data Analysis (TDA) identifies sTNFR1 among 5 preclinical trials as effective therapy after SCI. High-content data from 5 previous unpublished preclinical SCI trials were merged and topological data analysis was run in order to extract syndrome-level information. The multidimensional array of outcome measures was mapped to a data topology, with nodes (colored circles) representing clusters of subjects that share similar multivariate patterns, and edges (lines) that represent shared similarities between nodes. TDA on all animals in preclinical drug trials revealed a distinct separation of subjects based on injury severity (a). **b-e,** filtering the topology by different measures reveals how those with moderate injuries have a distinct, non-uniform pattern of recovery (**b)**. A robust recovery of forelimb function (black circles, b-c) not dependent on tissue pathology (**d-e**) was observed. Exploration of nodes of interest (black circle, e) revealed they were given the same 75 kdyne IH contusion injury (f), and that the robust forelimb recovery (**g**) was the result of a dose response to sTNFR1 treatment (**h)**. Heat maps: high-low for values (**a-e, g**), 100-0% for group membership(**f,h**). Principal components analysis on all hemicontusion data revealed a first principal component (PC1) that explained 27.1% of the total variance in the dataset (**i**). Hypothesis testing for an effect of sTNFR1 on PC1 revealed a significant dose response effect (polynomial contrast), indicating that sTNFR1 treatment has a multivariate effect that is detectable across the syndromic SCI space (**j**). Error bars represent standard error of the mean, * p < 0.05.

The beneficial effect of sTNFR1 was discovered by first mapping the multidimensional syndrome across the range of biomechanically-graded experimental cervical SCI severities in the VISION-SCI data commons^23,25^ (**Fig.1a-e**). Changing the filter to code for degree of recovery revealed a subset of uniformly moderate injury severities with unexpected gradation in functional recovery after cervical SCI. Changing the filter to depict variations in histopathological outcome measures revealed that the unusual recovery in these moderate injuries was not attributable to lesion size or white matter sparing. We then asked what conditions were associated with remarkable recovery, and discovered a correspondence between higher measures of behavioral improvement and the dose-response function from a blinded randomized preclinical trial of single bolus intrathecal sTNFR1 delivered to the cervical cord immediately after SCI unilateral contusion (**Fig. 1f**). Thus, in the heat map of the network shown in **Fig. 1h**, more animals from the vehicle group were located within the region of the network that matched poor forelimb recovery, while rats that received sTNFR1 i.t. were concentrated in nodes that had higher performance values in paw preference and grooming (**Fig. 1g**). No other similar relationships were found between drug treatment groups (minocycline, ciclopirox, methylprednisolone, DMSO) and recovery, although each of these had been identified in at least one study as showing efficacy in preclinical SCI^9,12,26^. The sTNFR1 effects appeared to be independent of gross measures of lesion size, white matter and gray matter sparing, or motor neuron numbers. Findings were confirmed by univariate analysis of each outcome; but importantly, none of those independent univariate analyses considered the variability of outcomes in the context of the whole ensemble of treatments and injury variables. Thus, TDA of the combined data from multiple studies appears to be useful in parsing out efficacy and identifying therapeutic promise in combined studies where multiple manipulations and outcomes can be examined in the aggregate.

The TDA showed a pattern suggesting that sTNFR1 had a unique effect on outcomes of combined variables, but did not provide a quantification of that effect. We therefore used the entire combined dataset of prior hemicontusion studies from our lab and applied principal component analysis (PCA) in order to evaluate effects of drug and the relative contribution of multivariables to each syndromic metric^27^. PC1 revealed significant loadings from multiple measures related to both neurological recovery and lesion size, accounting for 27.1% variance (**fig. 1i**). We then assigned z-scores for this PC for each rat in the sTNFR1 immediate delivery cohort and used analysis of variance to test for significant dose effects. PC1 showed a highly significant treatment effect, a quadratic contrast indicating an inverted U-shaped dose response function (ANOVA, p < 0.05; **Fig. 1j**). This analysis thus indicates that the aggregate syndromic metric PC1, derived from a large population of rats with SCI with and without drug treatment, was improved by sTNFR1 in a dose-dependent manner. Separate univariate analyses of neurological outcome measures, lesion volumes, and motor neuron sparing, revealed significant effects on forelimb function, but not on lesion measures (data not shown). These results suggest that immediate sTNFR1 i.t. treatment can affect long-term neurological outcomes.

Data-driven discovery of the sTNFR1 dose-dependent benefit was accomplished using a dataset with only limited lesion-related histopathological outcome measures, and neither the TDA nor the syndromic PCA detected a clear relationship between spared tissue and neurological function, perhaps implicating more subtle aspects of recovery (e.g. ‘plasticity’). While this result was promising, we recognized that the effects of immediate application needed to be extended to later time points if these results were to be relevant to developing therapies. Thorough exploration of time and dose-response variations with studies of long-term outcome is time and effort intensive, so we looked for early biological indices that might predict long-term outcome. We hypothesized that early neuroinflammation (NI) would be correlated with neurological outcomes, and early markers of NI could be used to test for acute effects of sTNFR1 that might translate into long term neurological improvements. Thus, we examined the effects of delayed (90 min postinjury) sTNFR1 on the early production of TNFα and related cytokines, as well as the time course of microglial activation and macrophage invasion over the first week after injury^1^ in a series of experiments. SCI produced a rapid increase in TNFα, IL1β and IL-6 protein at 3 hrs after injury, with return toward baseline levels at 24 hrs. Treatment with sTNFR1 decreased the level of TNFα at 3 hours, but did not significantly affect IL-1B or Il-6 (**Fig. 2a-c)**. Microglial/macrophage ‘activation’ followed, with a protracted development of ED1+ and Iba1+ staining intensity over the first week after SCI (**Fig. 2d, g, j and m**, ANOVA, sham vs. vehicle, p <0.05), as reported for other SCI models. Treatment with sTNFR1 90 minutes after injury significantly reduced ED-1 + expression at both 24 hours (**Fig. 2f,** ANOVA, vehicle vs. sTNFR1, p <0.05) and 7 days post injury (**Fig. 2i**, ANOVA, vehicle vs. sTNFR1, p < 0.05), and Iba1+ expression at 7 days post injury (**Fig. 2o**, ANOVA, p < 0.05).

**Figure 2.**
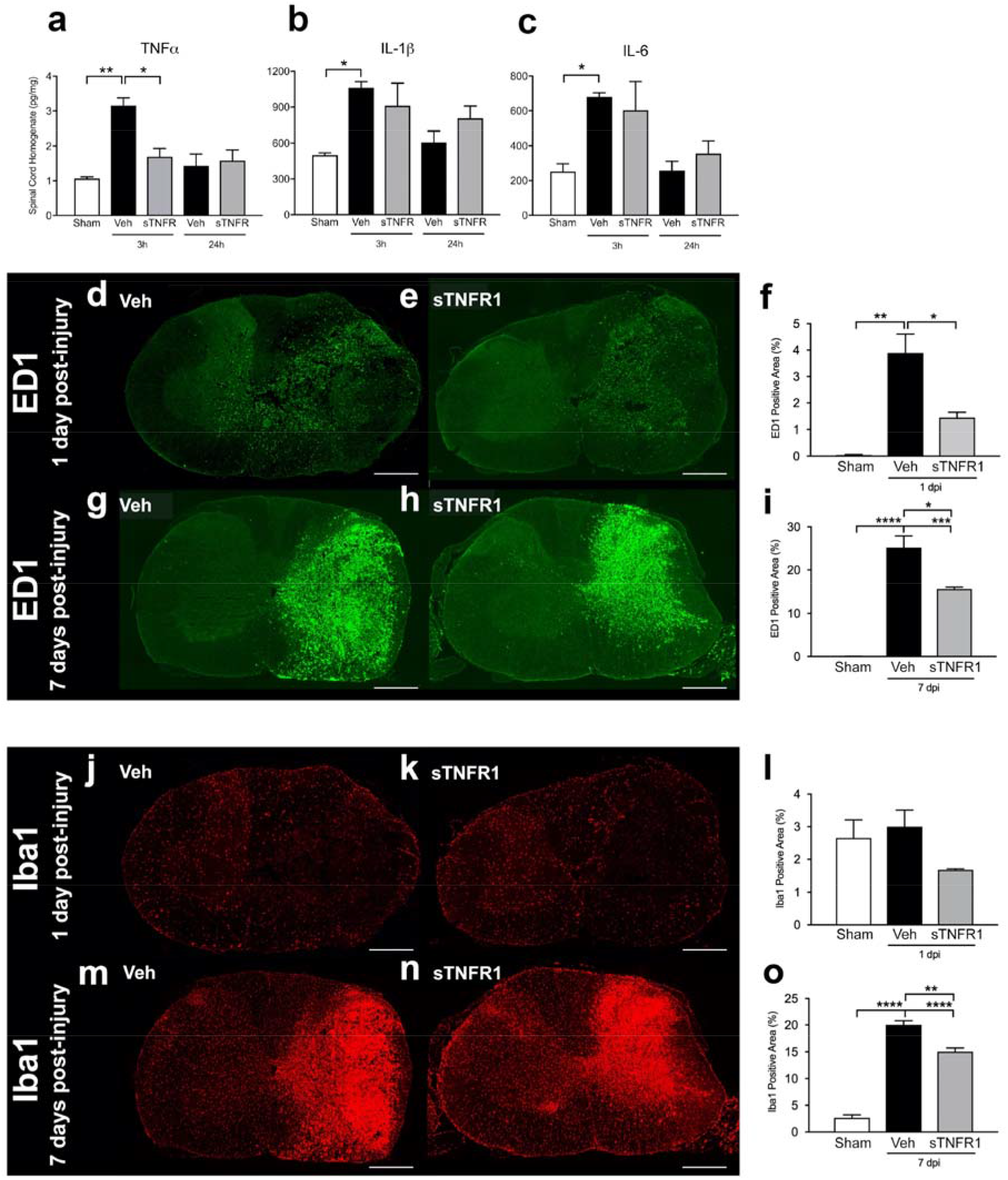
Acute sTNFR1 treatment reduces indicators of neuroinflammation. **a-c**, Spinal cord injury produces a strong early (3 hr) inflammatory cytokine (TNFα, IL1-β, and Il-6) protein expression. TNFα expression is significantly mitigated by sTNFR1 treatment. **d-e**, Immunohistochemical stain of injured spinal cord shows SCI-induced expression of ED1+ cells at 1 day and 7 days post-injury, and early i.t. sTNFR1 treatment significantly reduces ED1 at both 1 day and 7 days **(f,i). j-n**, Expression of Iba1+ cells following spinal cord injury at 1 day and 7 days post-injury. early i.t. sTNFR1 treatment did not affect Iba1 expression at 1 day post-injury, but did significantly reduce Iba1 expression by 7 days post-injury **(l,o).** Scale bars represent 400 um. Error bars represent standard error of the mean, One-way ANOVA, * p < 0.05, ** p < 0.01, *** p < 0.001, **** p < 0.0001.

Delayed sTNFR1 treatment at 90 minutes was chosen first because the literature and our preliminary data showed that TNFα was significantly increased at this time point, but returned to normal within several hours. Microarray PCR data showed substantial reversal of the inflammatory response at 3 hours as well (**Supplemental Figure 1**). Thus, the same dose of sTNFR1 that was effective when given immediately after SCI also had dramatic biological effects on neuroinflammation when delayed by 90 minutes.

We next used RNAseq to further examine the effect of sTNFR1 on the injured cord at the molecular level, using 3 groups of rats (Sham SCI, SCI+BSA Vehicle, SCI+sTNFR1, n=5 per group). Differential gene expression analysis revealed more than 5,000 genes induced upon SCI (Sham vs. BSA comparison) and 295 genes that were significantly up- or down-regulated after sTNFR1 injection (BSA vs. sTNFR1 comparison; adjusted p value < 0.05, **Fig. 3a**). A Gene Ontology enrichment analysis (**Fig. 3b**) showed that immune/inflammation-related functions were highly enriched as expected. Ninety-five of the 295 genes belong to the enriched inflammation-related gene ontologies (**Fig. 3b; Supplemental Table 2**). Next, we performed genome-wide gene co-expression network analysis, an unsupervised approach to identify groups of genes with similar expression patterns that capture a substantial amount of overall expression variation^28,29^. Each module was summarized by its first principal component (*co-expression module eigengene,* **Fig. 3c**), and the association strength of each gene with each module (*k*_ME_) was determined by correlating its expression pattern with each module eigengene.

**Figure 3.**
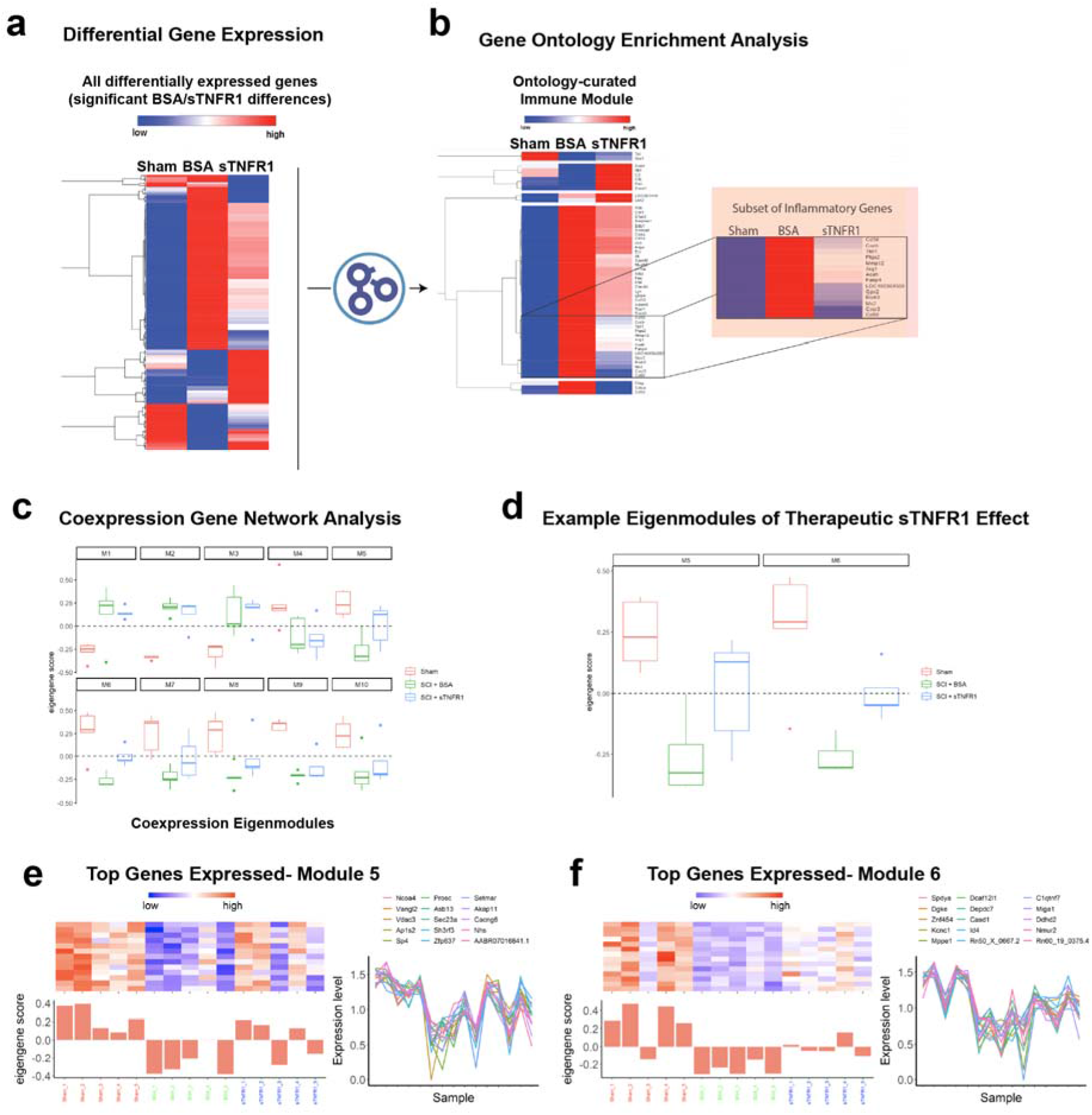
Differential gene expression in response to i.t. sTNFR1 treatment. RNAseq was used to explore the effect of sTNFR1 on the molecular level after SCI. **a,** We identified 295 differentially expressed genes (BSA vs. sTNFR1); 224 of those genes were altered by SCI (and BSA injection) and sTNFR1 administration reversed that change (partially or fully) to bring expression back to the pre-injury levels. **b**, Gene Ontology enrichment analysis was used to discover the processes that were most robustly affected by sTNFR1, revealing predominant inflammation/immune genes that were altered. A closer look at the ontology-curated immune module shows 52 genes to be involved in some capacity in immune and defense responses; sTNFR1 treatment reversed expression levels for a majority of these genes. **c,** Unsupervised gene coexpression network analysis identified 10 modules, which were summarized by their eigengenes and related to treatment effect**. d,** In two modules (M5, M6), sTNFR1 treatment significantly reversed the transcriptional phenotype compared to Sham and BSA vehicle treatment (one-way ANOVA, p < 0.05, error bars represent standard error of the mean). **e-f,** Module eigengenes and genes with the highest *k*_ME_ values (Pearson correlation to module eigengene) for modules M5 and M6, which best exemplified the therapeutic effect of sTNFR1.

Applying gene co-expression network analysis reduced the dimensionality of our dataset by more than 3 orders of magnitude and revealed 10 gene modules that were subsequently examined with respect to treatment condition (**Fig. 3c**). In 2 of these modules (arbitrarily designated M5 and M6), the SCI+Vehicle group changed its transcriptional phenotype in comparison to the Sham group, and upon sTNFR1 treatment a significant reversion of that phenotype was observed (ANOVA, p < 0.01; **Fig. 3d**). The eigengenes and top genes for these two modules, which best exemplified the therapeutic effect of sTNFR1, are illustrated in **Fig. 3e and f**. This transcriptomic approach confirms the effect of sTNFR1 treatment on SCI and highlights specific gene networks that are directly or indirectly regulated by TNFα and will be the main targets for intervention in future studies.

Finally, behavioral measures of gross and fine forelimb recovery were assessed over the course of 6 weeks post-SCI, as well as histological analysis of cell sparing, lesion size, and neuroinflammation markers. **Figure 4** shows the long-term outcomes from groups of rats receiving 90-min delayed sTNFR1 vs BSA vehicle controls. Each of the forelimb outcome measures was significantly better over 6 weeks in the sTNFR1-treated group (ANOVA, p < 0.05, **Fig. 4a-b**). There was a significant decrease in OX-42 staining 6 weeks after SCI in the sTNFR1-treated group compared to vehicle (ANOVA, p < 0.05, **Fig. 4c-d**). Therefore, the degree of persistent neuroinflammation may be a factor in determining behavioral outcome, and acute bolus treatment with sTNFR1 that affects early TNFα production/expression appears to have a lasting effect on the development of neuroinflammation and on neurobehavioral outcomes^30^. Mironets et al (2018) have provided additional evidence for a critical role for sTNF signaling in the production of long term neurological and immune dysfunction after thoracic SCI^31^. Thus, multiple approaches and experiments lead to the conclusion that modulation of the sTNF signaling offers a viable approach to therapeutic treatments for SCI. Accordingly, the advanced TDA and transcriptomics analyses we employed to identify sTNFR1 as an effective delayed treatment for cervical SCI may be useful in streamlining the evaluation of the pharmacological treatments aimed at improving post CNS recovery of function. In addition, the ability to identify novel gene expression modules that associate with better outcomes is likely to be useful for identifying novel therapeutic targets that could be tested in a variety of preclinical neurotrauma models.

**Figure 4.**
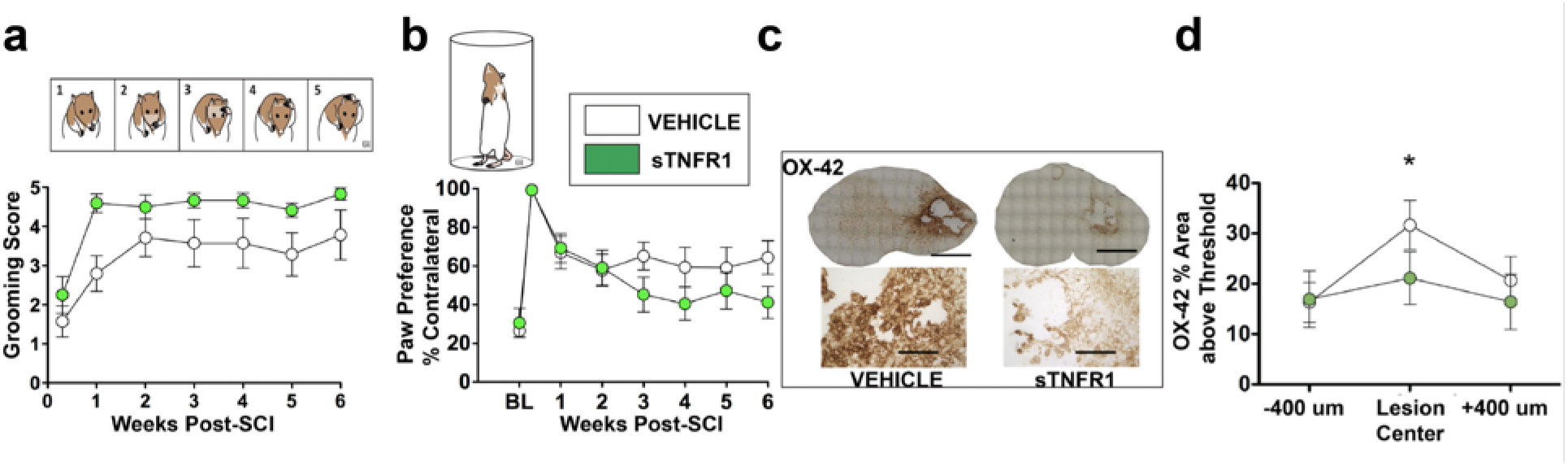
Acute sTNFR1 treatment improves behavioral recovery and mitigates chronic neuroinflammation. Tests of forelimb recovery following unilateral cervical contusion were assessed for 6 weeks post-injury. Subjects that received a single acute i.t. sTNFR1 bolus showed significantly better scores on grooming (**a**) and paw placement (**b**) tests over 6 weeks (Repeated-measures ANOVA, Time x Treatment, p < 0.05). Terminal histological assessment of neuroinflammation at 6 weeks post-injury (**c)** show that sTNFR1 treatment significantly reduced OX-42 expression (**d,** ANOVA, p < 0.05) indicating that early intervention with sTNFR1 was sufficient to block the development of lasting neuroinflammation. Error bars represent standard error of the mean, * p < 0.05.

## Acknowledgements

We would like to thank (in alphabetical order) Tomoo Inoue, Yvette Nout Ellen Stuck, and Jason Talbott for useful comments on a prior version of this manuscript. This work was funded by NIH grants NS067092 (A.R.F.), NS069537 (A.R.F), NS038079 (J.C.B. and M.S.B.), AG032518 (M.S.B. and J.C.B.), NYSCoRE CO19772 (M.S.B. and J.C.B.), and NIH UCSF Anesthesia T32 training grant (NIGMS T32GM008440) to JZP.

## ONLINE METHODS

### Animals

Female Long-Evans hooded rats (Simonsen Laboratories, Gilroy, CA, USA) with a mean weight of 230 g and mean age of 77 days were used in this study. Rats were triple-housed in plastic cages, maintained on a 12-hour light/dark cycle, and had access to food and water *ad libitum*. All experiments were approved by the Institutional Laboratory Animal Care and Use Committee of the University of California at San Francisco and were performed in compliance with NIH guidelines and recommendations. Surgical procedures were carried out under aseptic conditions, during which animals were kept under deep anesthesia induced and maintained by isoflurane inhalation (IsoFlow, Abbott Laboratories, North Chicago, IL, USA; 2-3%). Anesthetic plane was monitored by testing withdrawal to foot pinch. Cefazolin (Ancef, Novation, LCC, Irving, TX) 25 mg/kg, was administered prior to surgery and for 3 days postoperatively (for chronic injured subjects). Lacrilube ophthalmic ointment (Allergan Pharmaceuticals, Irvine, CA, USA) was applied to the eyes prior to surgery and body temperature was monitored using a rectal thermal probe and maintained at 37.5 + 0.5 C using a heating pad.

### Unilateral cervical spinal contusion injury

Spinal cord injury was delivered as described previously^30^. Briefly, a dorsal, midline skin incision was made, the skin dissected, and the trapezius muscle was cut just lateral to the midline from C1/2 to T2. Muscle layers were dissected to expose C3-T1 spinous processes. A dorsal laminectomy was then performed at C5 to expose the spinal cord. Contusion injury was produced using an Infinite Horizons Impactor (Precision Systems, Fairfax, VA, USA) fitted with a 2mm impactor tip that was centered over the right side of the C5 spinal segment. Impactor was computer-controlled to consistently impact the spinal cord at a force of 75 kilodynes. After injury, trapezius muscle was sutured and skin incision closed with wound clips. The analgesic buprenorphine (0.05 mg/kg), and the antibiotic Cefazolin (50mg/kg, Henry Schein, Melville, NY) were administered, and the animal recovered overnight in an incubator (Thermocare Intensive Care Unit with Dome Cover; Thermocare, Inclined Village, NV). All animals were inspected daily for wound healing, weight loss, dehydration, autophagia and discomfort. Appropriate veterinary care was provided when needed.

### Drug Delivery

Human recombinant soluble TNF receptor 1 (sTNFαR1; R&D systems) was delivered by way of an intrathecal cannula (30 cm, PE-10, sterilized with 95% ethanol and prefilled with sterile 0.2% BSA vehicle) that was placed in the cisterna magna and threaded 1 cm caudally into the subarachnoid space until it was visible under the dura within the laminectomy site at C4/5, and then positioned so that it was directly rostral to the laminectomy window. The cannula tip was positioned over the hemicord, contralateral to the target site for unilateral contusion. The cannula was fixed in place using two bilateral finger trap sutures into the sternohyoid muscles. Subjects were then placed in the Inifinite Horizons impact device and the external cannula tip was heat flanged and then connected to a 10 μl Hamilton syringe filled with sterile sTNFR1 solution or vehicle in a blinded, randomized fashion. Immediately following contusion injury, drug was delivered intrathecally over 5 minutes followed by a 20μL vehicle (sterile, 0.2% BSA) flush over 10 minutes. Following drug delivery, finger trap sutures were removed, the cannula was carefully withdrawn, and the surgery site was closed using standard surgical procedures^32,33^. Subjects were carefully monitored post-operatively for signs of bilateral deficits, indicating potential damage by the cannulization procedures. Ciclopirox was delivered subcutaneously at a concentration of 10 mg/kg. The vehicle control for the ciclopirox study was dimethyl sulfoxide (DMSO). DMSO was delivered subcutaneously at a concentration of 1 mg/kg. Methylprednisolone was given intravenously at a concentration of 30 mg/kg. Minocycline was delivered intraperitoneally at a concentration of 45 mg/kg.

### Paw Preference Test

Animals were placed in a clear plastic cylinder that is situated in mirrored corner, so that the animal can be viewed from all angles. Animals are filmed on a digital camera while they explore the cylinder for 3 minutes. Slow motion high-definition playback of each session allows a rater who is blind to condition to record each time a weight-supported placement is made on the cylinder wall by either the left forepaw alone, right forepaw alone, or both. A “left” or “right” count was given if the other limb did not contact the side of the cylinder within 0.5 sec of the initial placement. A “both” count was given if both forepaws were placed on the cylinder within 0.5 sec of each other. During lateral exploration, a “both” score was also given for each two step “walking” sequence, during which both paws changed position on the cylinder wall. If one paw remained in place while the other was placed on different parts of the cylinder, a count was not given until the anchored paw was lifted.

### Grooming Test

An assessment of stereotypical grooming behavior was adapted for use in cervical model of SCI by Gensel et al., and used to determine recovery of forelimb range of motion^32^. Cool tap water was applied with gauze to the animal’s head and back, and then the animal was placed in a clear plastic cylinder with mirrors on either side so that the animal could be observed from all angles. Grooming activity was recorded with a digital video camera from the onset of grooming through at least 2 grooming sequences. Slow motion high-definition playback was used to score each forelimb independently, using a 6-point scoring system as follows: 0 indicates the animal is unable to make contact with the forepaw to any part of the face or head; 1 indicates the animal’s forepaw can make contact with the underside of the chin and/or mouth area; 2 indicates the animal’s forepaw can make contact with the area between the nose and eyes, but not the eyes; 3 indicates the animal’s forepaw can make contact with the eyes and the area up to, but not including, the front of the ears; 4 indicates the animal’s forepaw can make contact with the ears, but not the area of the head behind the ears; 5 indicates the animal’s forepaw can make contact with the area of the head behind the ears. Animals were tested at 2, 7, 14, 21, 28, 35, and 42 days after injury.

### Irvine, Beatties, Bresnahan (IBB) forelimb rating scale

Fine forelimb function was assessed using a cereal eating test as described in Irvine et al.^33^. Animals were individually placed in their home cages and given doughnut- and spherical-shaped pieces of cereal, and eating was filmed with a digital video camera. Slow motion high-definition playback was used to evaluate forepaw use. Evaluation was made using a standardized scoring of forelimb behaviors while eating (e.g. joint position, object support, digit movement, and grasping technique). An IBB score was assigned using the 10 point (0-9) ordinal scale for each shape.

### Histological preparation

Animals were perfused through the left ventricle of the heart with 4% paraformaldehyde under deep anesthesia with pentobarbital. The cords were removed and postfixed in 4% paraformaldehyde for 2 hr and then cryoprotected in PBS containing 30% sucrose. A 2 mm block containing the lesion epicenter was then incubated in 100% OCT for 2 hr and then mounted in a cryomold (filled with OCT) in coronal orientation and rapidly frozen using dry ice. The blocks were stored at −80°C until sectioning. The cords were cut coronally at 20 um and every section was retained and mounted. Sections were stained with Luxol fast blue for myelin/white matter integrity and counterstained with Cresyl violet or for cell body assessment.

### Sparing at lesion epicenter

A camera lucida drawing of the section with the largest extent of lesion (the lesion epicenter) was made outlining intact gray and white matter, and the lesion. Pixel counts from digitized drawings in Adobe Photoshop 5.5 (Adobe Systems Inc., San Jose, CA) were used to determine the area of spared tissue for both hemi-cords at the lesion center. The percent sparing for the ipsilateral hemi-cord was determined by dividing the total spared ipsilateral tissue area, spared white matter tissue area, or spared gray matter tissue area, by the same measure from the contralateral hemi-cord [(ipsilateral spared tissue area/contralateral spared tissue area)x100]. This normalized within subjects and corrected for any biological differences in spinal cord size or tissue preparation.

### Immunohistochemistry

Fixed spinal tissue sections were blocked and permeabilized for 1 h with 10% normal donkey serum and 0.3% Triton X-100. The sections were then incubated overnight at room temperature (RT) with mouse monoclonal antibody for ED1 (1:300; Bio-Rad Laboratories, Hercules, CA, USA) and Iba-1 (1:500; Fujifilm Wako Chemical. Richmond, VA, USA). After washing with phosphate-buffered saline (PBS) 2 ml, the slides were incubated for 1 h at RT with fluorescent (Alexa 488 and 594) donkey anti-mouse secondary antibody (1:1000; Life Technologies, Carlsbad, CA, USA). The slides were briefly rinsed with PBS 2 ml and coverslipped with VECTASHIELD ® containing 40,6-diamidino-2-phenylindole (Vector Laboratories, Burlingame, CA, USA). The stained spinal tissue sections were photographed using the BioRevo fluorescence microscope BZ-9000 Generation II (Keyence, Itasca, IL, USA). Fluorescence was measured using BZ-9000 Generation II analyzer (Keyence) and analyzed by NIH ImageJ. Proportional area measurements were acquired by adjusting the thresholds of stained sections in image J and ratio of immunoreactive area to the section area was calculated. Every eighth section of the spinal cord was analyzed for a distance of up to 5 mm in the rostral and caudal directions from lesion center.

### OX-42 staining

Slides were first washed 2 x 5 min in Tris-buffered phosphate (TBS) with Triton X-100 (0.025%). Slides were then blocked in 10% normal goat serum with 1% bovine serum albumin (BSA) in TBS for 2 h at room temperature. Primary antibody (anti-OX-42, 1:500, Abcam) was then added, and incubated overnight at 4° C. Slides were then rinsed 2 x 5 min in TBS+ Triton X-100, and incubated in horseradish peroxidase (H_2_O_2_) in TBS with 1% BSA for 15 min. To visualize reaction, DAB chromagen with nickel enhancement was added, followed by dehydration and tissue clearing.

### OX-42 quantification

As specific microglia cell counting is often obfuscated by clustered and overlapping cells, quantification of OX-42 expression was performed by calculating area of positive staining on the injured hemisphere, and correcting by the contralateral uninjured hemisphere. Threshold for positive staining was determined using MetaMorph image analysis software. Briefly, an image analysis macro script was written to first set the relative white balance for each image, then determine the ratio between number of pixels above threshold and the total number of pixels in the area. This process was repeated for both ipsi- and contralateral sides of the spinal cord, and the final percentage of OX-42 expression was formulated by dividing pixels above threshold on ipsilateral side by pixels above threshold on contralateral side.

### Luminex multiplex cytokine assays

Total cellular proteins were prepared from spinal cord tissue. Aliquots were analyzed for inflammatory cytokines, with Luminex xMAP multiplexing technology (Luminex Corp., Austin, TX). Spinal cord protein specimens were prepared for analysis in a 96-well plate utilizing a custom 7-cytokine Milliplex MAP Rat Cytokine/Chemokine Magnetic Bead Panel (RECYTMAG-65K, Millipore Corp., Billerica, MA) following the kit-specific protocols provided by Millipore. Analytes were quantified using a Magpix analytical test instrument, which utilizes xMAP technology (Luminex Corp., Austin, TX), and xPONENT 4.2 software (Luminex Corp.). xMAP technology uses fluorescent-coded magnetic microspheres coated with analyte-specific capture antibodies to simultaneously measure multiple analytes in a specimen. After microspheres have captured the analytes, a biotinylated detection antibody binds to that complex. Streptavidin PE then attaches as a reporter molecule. Inside the instrument, magnetic beads are held in a monolayer by a magnet, where two LEDs are used to excite the internal microsphere dye and the dye of the reporter molecule, respectively. A CCD camera captures these images, which are then analyzed by Milliplex Analyst software (Millipore).

### Assay of inflammatory chemokine and cytokine gene expression

Acute injured animals (3 hours post-injury) were deeply anesthetized with 5% isoflurane. Animals were given a brief intracardiac saline perfusion, and then a section of spinal cord (centered rostrocaudally around the lesion) were removed, flash-frozen in isopentane, and then stored at −80 C. To assess changes within and between subjects, cords were split on the midline, to produce injured and uninjured hemicords. RNA from spinal hemicords were extracted and purified using an RNEasy Lipid Minikit (Qiagen) and RNA concentration was quantified using a Biomate 3 spectrophotometer (Thermo Scientific) set to read absorbance at 260 nanometers. Equal amounts of RNA per hemicord were subjected to reverse transcriptase reaction (RT2 First Strand Kit, SABiosceinces) to yield cDNA. Equal volumes of cDNA for each hemicord were pooled by condition, and subjected to PCR amplification using a commercially available multitarget Inflammatory cytokine/chemokine array (Qiagen). One array was used per pooled condition. PCR data were then normalized to housekeeping genes and fold changes in mRNA expression between conditions were calculated using the Delta-Delta Ct Method^34^.

### RNA extraction

1 cm of hemicord was homogenized in 1 ml of TRIZOL solution (Thermo Fischer #15596018) and subsequently total RNA was extracted using the manufacturer’s protocol. RNA was purified one additional time using 3M sodium acetate.

### Library preparation

1 μg of extracted RNA was used for the library synthesis. DNA library was synthesized using Illumina’s TruSeq Stranded Total RNA with Ribo-Zero Globin and by following the manufacturer’s instructions. Libraries were then quantified and tested for proper fragmentation using the 2100 Agilent bioanalyzer and the Agilent DNA 1000 kit (Agilent # 5067-1504).

### RNAseq

Samples were sequenced on Illumina’s HiSeq 4000 aiming for 40 million single-ended reads per sample.

### Bioinformatic analysis

We used the software packages Scythe and Sickle to trim reads as necessary to achieve maximum quality. We aligned the trimmed reads against the rat genome and transcriptome using the TopHat2/Bowtie2 software. After read alignment, we used the featureCount function in R to get the raw counts of every transcript in our samples. Data were examined for outliers using the SampleNetwork R function^35^ and batch effects were corrected using the ComBat function^36^ of the sva package in R^37^. Unsupervised gene co-expression network analysis^28,29^ was performed using a four-step approach. First, pairwise biweight midcorrelations (bicor) were calculated for all analyzed transcripts over all samples using the bicor function in the WGCNA R package^38^. Second, transcripts were clustered using the flashClust implementation of a hierarchical clustering procedure with complete linkage (minimum cluster size = 12) and 1 – bicor as a distance measure. Third, the resulting dendrogram was cut at a height corresponding to the top 10% of pairwise correlations. Fourth, modules were summarized by their eigengenes^39^, defined as the first principal component obtained by singular-value decomposition of the coexpression module, and highly similar modules were merged if the correlations of their module eigengenes exceeded 0.85. This procedure was performed iteratively, such that the pair of modules with the highest correlation (>0.85) was merged first, followed by recalculation of module eigengenes, followed by recalculation of all correlations, until no pairs of modules exceeded the threshold. The WGCNA measure of intramodular connectivity (*k*_ME_) was then calculated for each gene with respect to all modules by correlating its expression pattern across all samples with each module eigengene.

### Statistical Analyses

Topological data analysis (TDA) was conducted according to methods previously described^2^. Briefly, TDA uses an ensemble machine learning algorithm to rapidly iterate across subject bins, resampling the metric space and replacing subjects after each sampling. This resampling procedure ultimately arrives at the most stable ‘consensus vote’ that best represents the multidimensional data shape. The result of this approach is a clustering of subjects (‘nodes’) and the connections between these clusters (‘edges’ or lines). Variables and outcomes of interest that were used in the TDA included injury condition, drug condition, behavioral endpoints, and histological endpoints (tissue sparing, lesion size, etc). Heat maps for the color schemes of the flares represent the range of highest values (red) to lowest values (blue) for each respective outcome being visualized.

Principal component analysis was performed by eigenvalue decomposition of the correlation matrix of all outcomes in using the FACTOR subcommand in SPSSv.18. This method extracts PCs that are uncorrelated (orthogonal) partitions of the variance. PCs were retained using 3 criteria: 1) retaining PCs with eigenvalues > 1.0 (Kaiser rule) ^40^ 2) Scree plot^41^, and 3) the over-determination of the factors^42^, retaining factors with at least 3 loadings above |.4|. PCs meeting all three criteria were examined and named using loadings above |.4|, thereby accounting for at least 20% of the variance. Visualization of PC loadings was achieved using the syndRomics package in the R statistical software program. To test for univariate effects of drug treatment on principal component scores, protein expression, behavior, and eigenmodule scores, we performed analyses of variance (ANOVA) using the SPSS GLM subcommand or R statistical software. Significant ANOVAs were followed by Tukey’s posthocs. To assess dose response we performed an additional polynomial contrast analysis. Significance for all univariate effects was assessed at p < .05.

---

**Supplemental Table 1.** Anti-inflammatory drug studies. Parameters of drug type and dose for each drug study that was included in the initial data-driven analysis of multivariate treatment effects.

| Study | Drug Delivery | Dose | SCI Model | SCI Severity | SCI Level | Laterality | Number of animals |
| --- | --- | --- | --- | --- | --- | --- | --- |
| Methylprednisolone | intravenous | 30mg/kg | NYU Impactor | 12.5mm drop | C5 | Unilateral | 10 |
| Minocycline | intraperitoneal | 45mg/kg | NYU Impactor | 12.5mm drop | C5 | Unilateral | 11 |
| DMSO | subcutaneous | 1mg/kg | IH Impactor | 100 kilodyne | C5 | Unilateral | 12 |
| Ciclopirox | subcutaneous | 10mg/kg | IH Impactor | 100 kilodyne | C5 | Unilateral | 8 |
| sTNFR1 | intrathecal | 0.035-3.5ng | IH Impactor | 75 kilodyne | C5 | Unilateral | 32 |
| Bovine Serum Albumin vehicle | intrathecal | 0.002 | IH Impactor | 75 kilodyne | C5 | Unilateral | 17 |

**Supplemental Figure 1.**
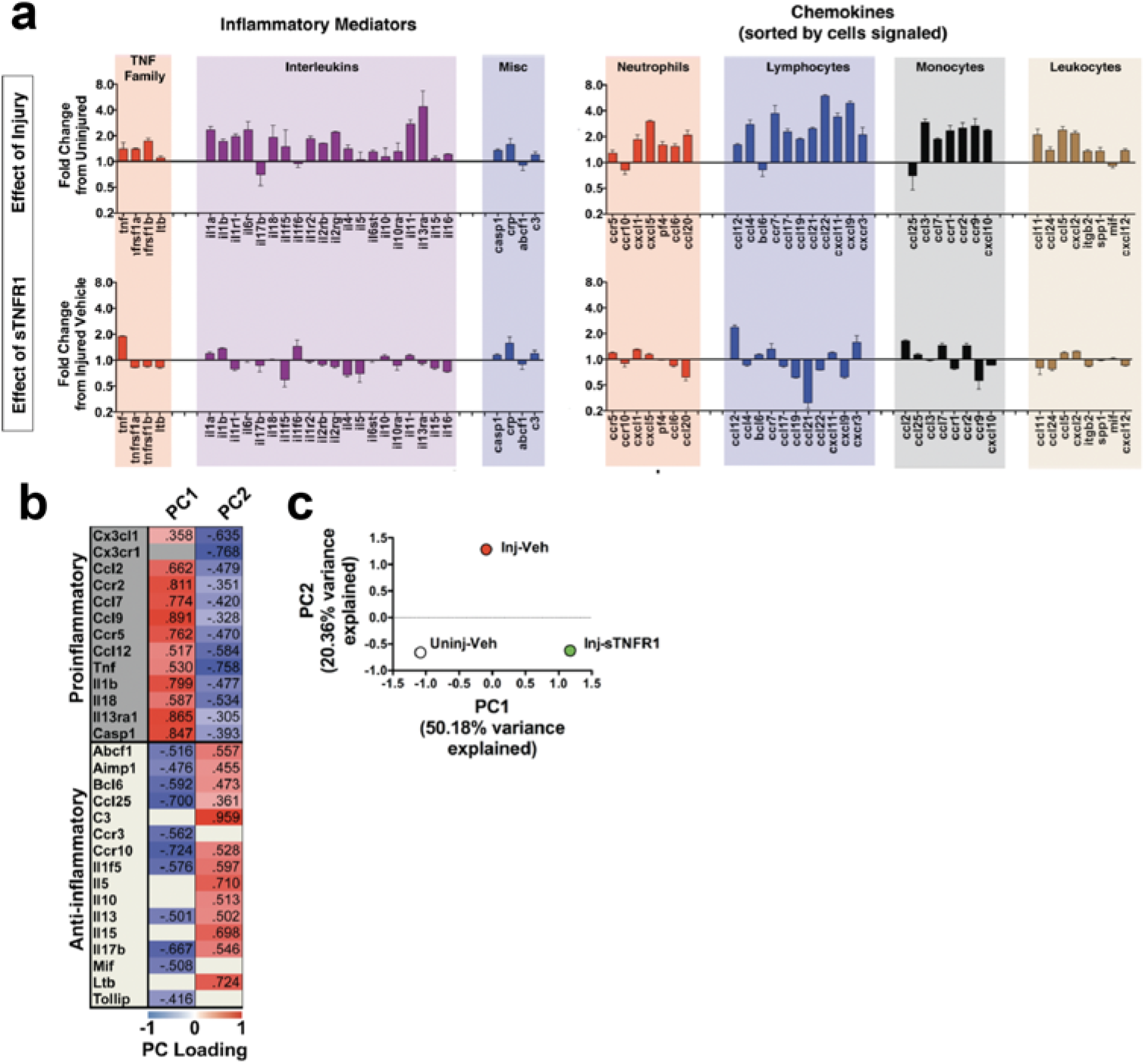
Effect of sTNFR1 on inflammatory gene panel. A multiplexed PCR panel of 84 inflammatory genes was used to assess the broad effect of sTNFR1 treatment on neuroinflammation after SCI. sTNFR1 was delivered i.t. 90 minutes after injury, and spinal cord was harvested for PCR 3 hours after injury. **a**, Overall expression levels were increased in the vehicle-treated SCI group, as expressed by fold-change relative to uninjured sham control subjects (“Effect of Injury”). sTNFR1 treatment either dampened or reversed this neuroinflammatory response in the majority of genes tested. (“Effect of sTNFR1”). **b**, Principal components analysis of all gene expression values across all testing groups revealed two principal components that together accounted for 71% of the variance. PC1 was characterized by high positive loadings in classically pro-inflammatory genes, and PC2 was most strongly driven by anti-inflammatory genes. **c,** biplot of the three experimental groups on the PC1 and PC2 axes illustrates the 2 dimensional syndromic space occupied by each experimental condition. sTNFR1 treatment (Inj-sTNFR1) produces an inflammatory profile that is distinct from both injury alone (Inj-Veh) and uninjured controls (Uninj-Veh).

**Supplemental Table 2.**
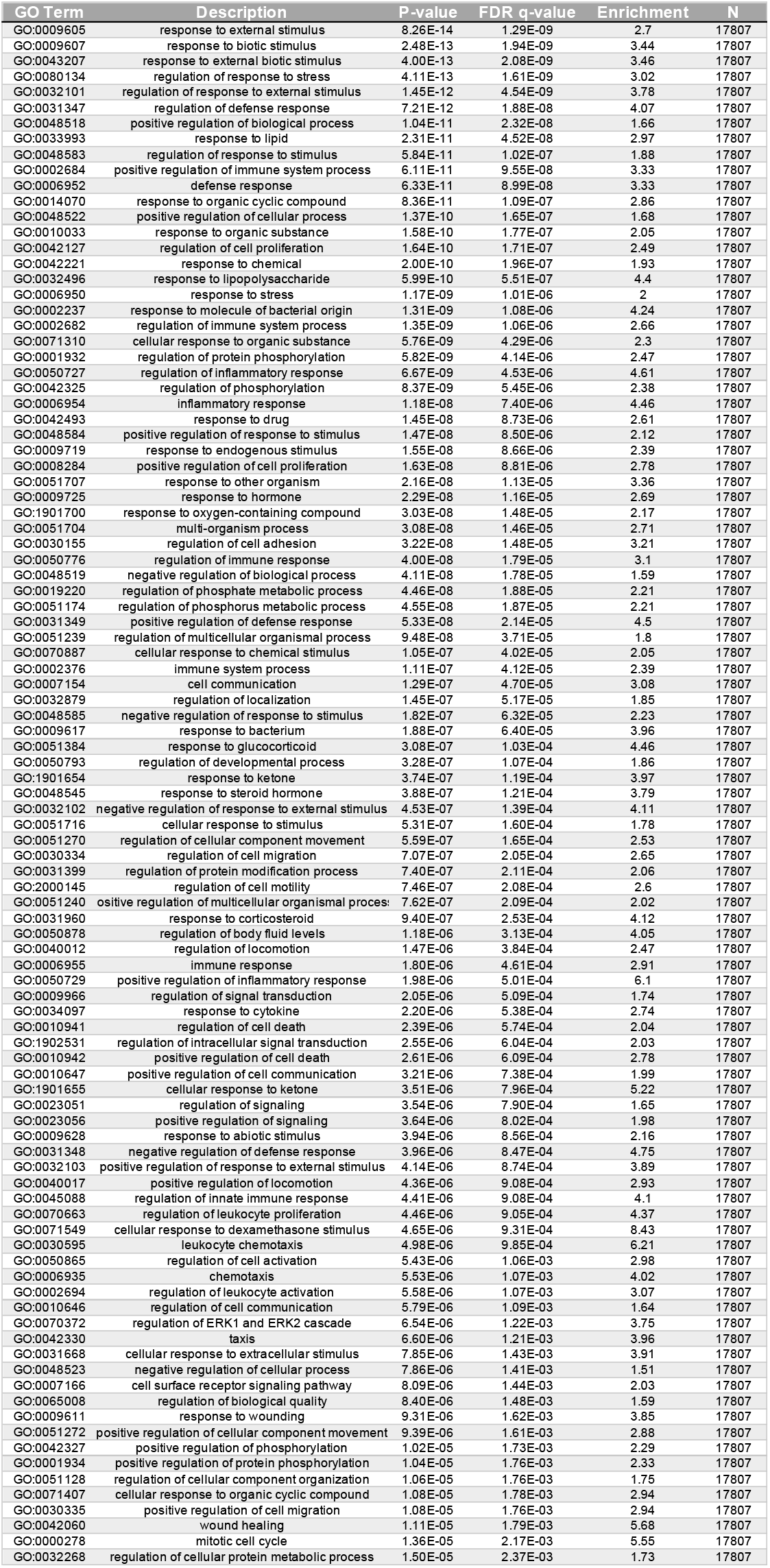
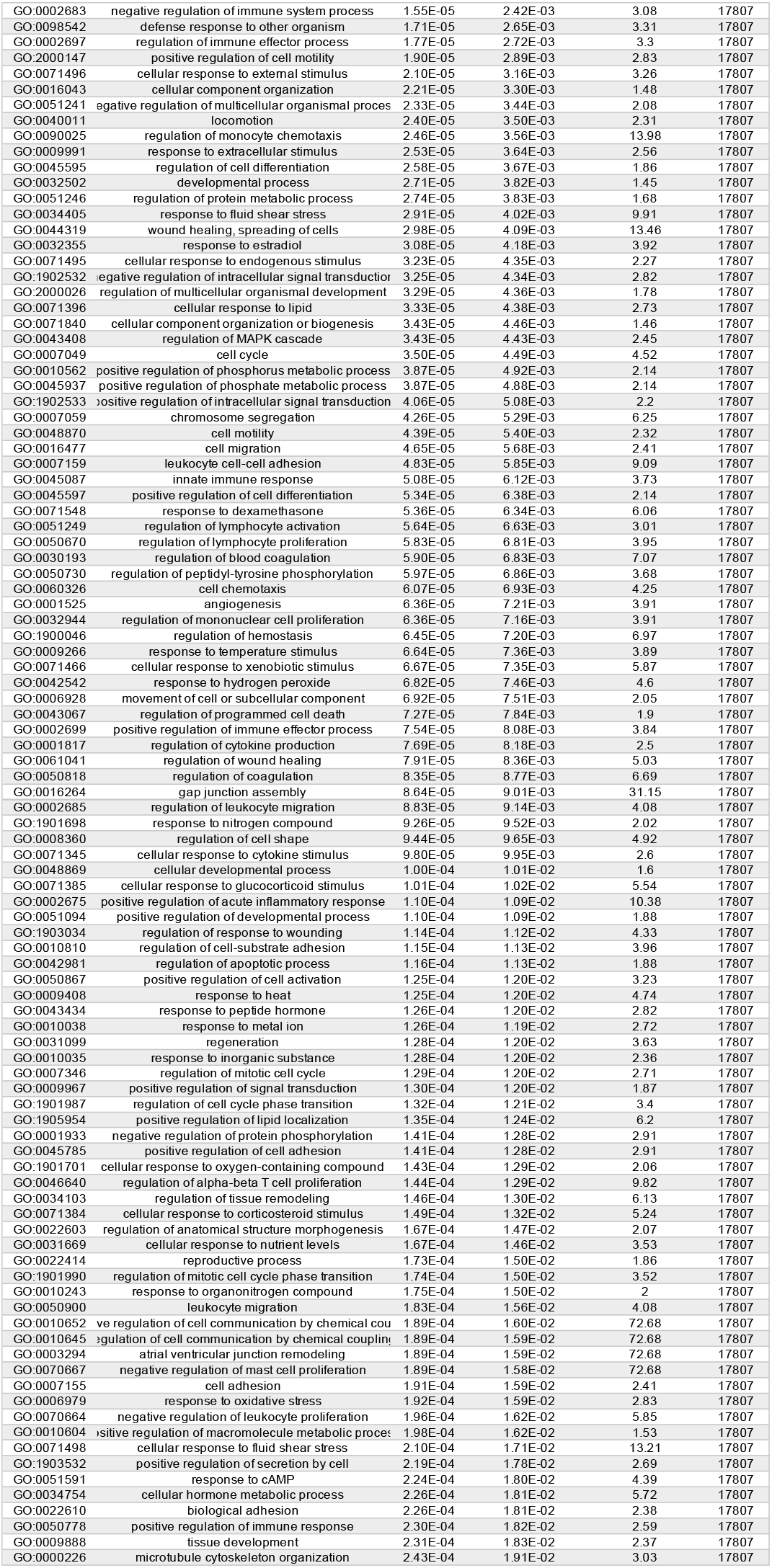
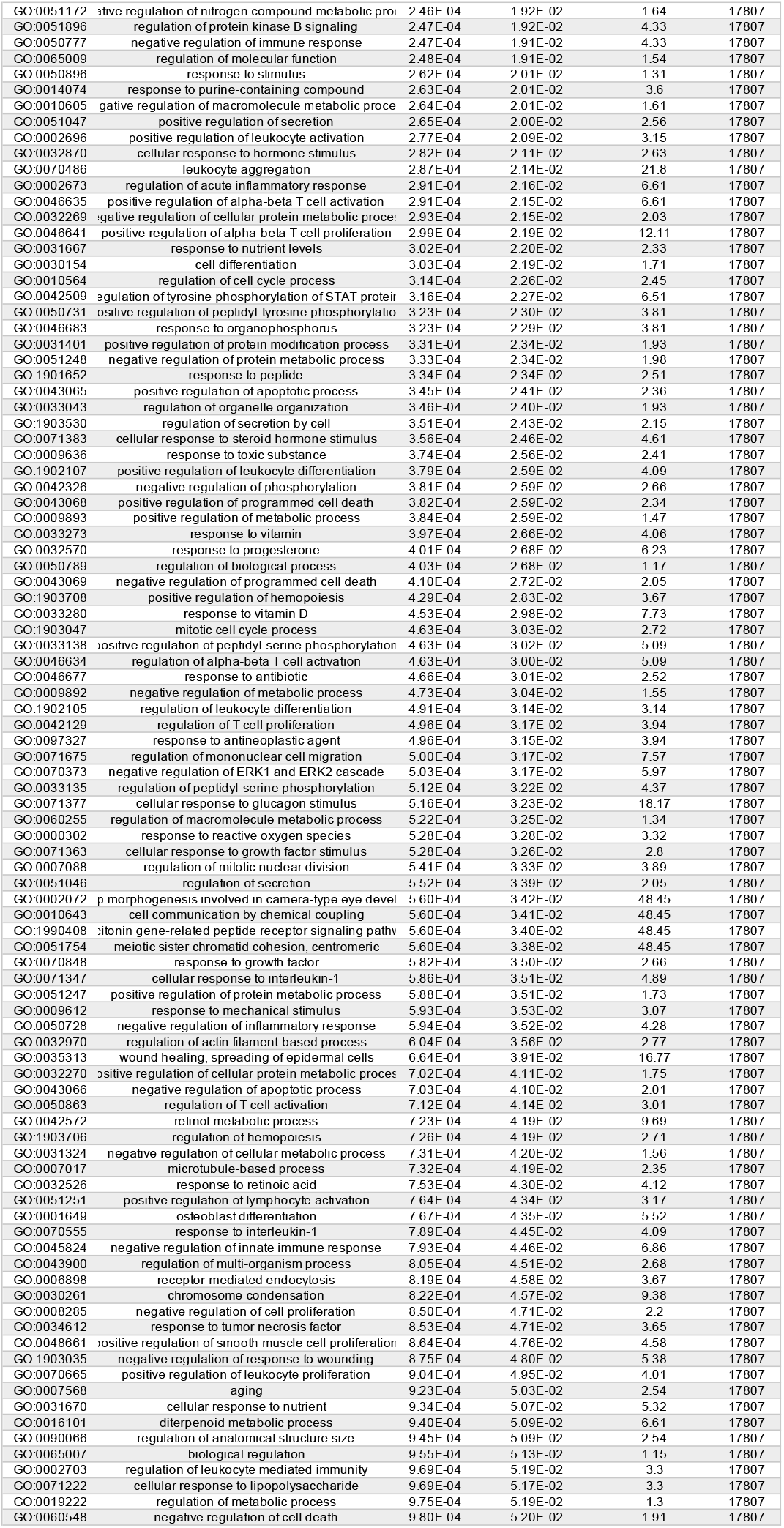
295 genes were significantly up- or downregulated after sTNFR1 injection. Terms from Gene Ontology analysis indicate that 95 of 295 (32.2%) differentially expressed genes were related to inflammatory processes. Columns 1 and 2 indicate the specific Gene Ontology term numbers and descriptions. P-value indicates nominal significance value for each gene, while FDR q-value reflects the adjusted value accounting for false discovery rate. Enrichment score indicates the degree to which the genes are overrepresented at the top or bottom of the entire ranked list of genes.

